# A Mathematical Modeling Approach to The Cort-Fitness Hypothesis

**DOI:** 10.1101/506741

**Authors:** Fadoua El Moustaid, Samuel J. Lane, Ignacio T. Moore, Leah R. Johnson

## Abstract

The Cort-Fitness hypothesis has generated much interest from investigators integrating field endocrinology with evolutionary biology, ecology, and conservation. The hypothesis was developed on the assumption that if glucocorticoid levels increase with environmental challenges and fitness decreases with environmental challenges, then there should be a negative relationship between glucocorticoid levels and fitness. However, studies across diverse taxa have found that the relationship between glucocorticoid levels and fitness is not consistent: some studies show a positive relationship, others negative, and some show no correlation. Hence, support for the hypothesis is not consistent and thus a deeper understanding of the mechanisms underlying the relationship between glucocorticoid levels, environmental pressures, and fitness is needed. We propose a mathematical model representing the links between glucocorticoid levels, environmental challenges, and fitness. Our model explores how variation in the predictability and intensity of environmental challenges, reproductive strategies, and fitness metrics can all contribute to the variability observed in empirical tests of the Cort-Fitness hypothesis. We provide qualitative results showing the Cort-Fitness relationship for different environmental scenarios and discuss how the model can be used to inform future Cort-Fitness studies.

## 1 Introduction

Organisms face differing levels of abiotic and biotic challenges throughout their lifetimes that can dramatically affect their fitness (Ricklefs and Wikelski, 2002; Lanctot et al., 2003; Satterthwaite et al., 2010). To survive and reproduce, animals must respond to these challenges through behavioral strategies, physiological responses, or a combination of both (Sapolsky et al., 2000). For vertebrates, the physiological response may involve changes in hormone levels and/or other regulating factors, often in an effort to maintain homeostasis (Sapolsky et al., 2000; Tilgar et al., 2017). Researchers often use these changes in circulating hormone levels as an indication of the conditions individuals are experiencing (Sapolsky et al., 2000; Romero and Wikelski, 2001). For instance, levels of corticosterone (cort), the predominant glucocorticoid in birds, have been used as a measure of environmental challenges that the animals are facing (Satterthwaite et al., 2010; Grunst et al., 2014; Kitaysky et al., 2001).

As glucocorticoids are metabolic hormones as well as stress hormones, elevated plasma levels have often been considered an indication that an individual is in poor body condition which is often associated with lower quality habitat (Romero and Wikelski, 2001; Bonier et al., 2009a; Kitaysky et al., 2003; Jaatinen et al., 2013). These relationships between plasma glucocorticoid levels and body condition have led to the assumption that individuals or populations with elevated levels of the hormone will exhibit decreased fitness, either through decreased survivorship, decreased reproductive success, or both. The Cort-Fitness hypothesis, first proposed in the early 2000’s and then formalized by Bonier et al. (2009a), tests the premise that if baseline glucocorticod levels increase with environmental challenges, and fitness decreases with environmental challenges, then baseline glucocorticoid levels and fitness should be negatively related (see Figure 1) (Sapolsky et al., 2000; Bonier et al., 2009a; Escribano-Avila et al., 2013). However, support for the hypothesis was not consistent (Bonier 20019a). Indeed, emerging evidence suggests that the effects of environmental challenges are more complicated, and individuals with slightly elevated glucocorticoid levels may have higher fitness than individuals with either low or significantly higher levels of glucocorticoids (Bryan et al., 2014; Bonier and Martin, 2016). In other words, the relationship between baseline glucocorticoid levels and fitness can be positive, negative, non-linear, or there may be no relationship at all. To date, the Cort-Fitness hypothesis has been tested primarily through observational studies (Bonier et al., 2009a; Bryan et al., 2014; Breuner et al., 2008; Crossin et al., 2012; Hood et al., 1998; McAuley et al., 2009). Here we use a mathematical model to test the effect of different scenarios of environmental challenges and glucocorticoid levels on an individual’s fitness in an effort to better understand the functional nature of the relationships between stressor, glucocorticoid levels, and fitness. Although we use birds as the inspiration for the model and calibrate model parameters accordingly, the model is quite general and adjustable to other study systems. With our model, we aim to address the following questions:

1. How may glucocorticoid levels vary with environmental challenges?
2. How may individuals life history stage affect their fitness?
3. How may individuals short-term response to stress affect the long-term trade off strategy?

**Figure 1:**
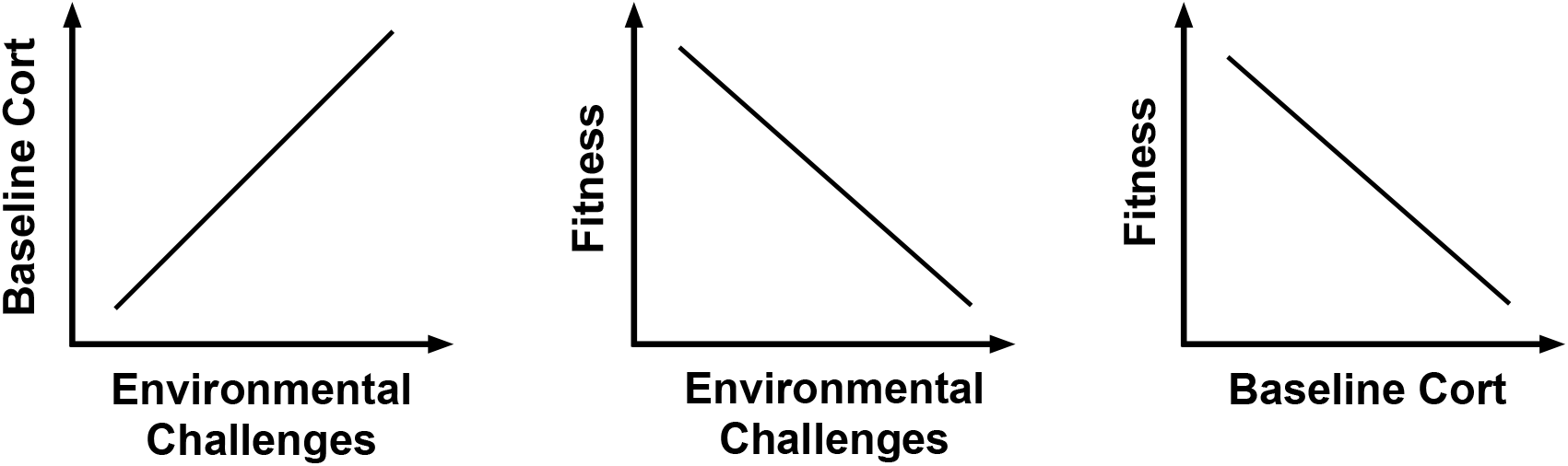
The Cort-Fitness Hypothesis schematic description from (Bonier et al., 2009a) (reprinted with permission). Baseline glucocorticoids increases with environmental challenges and individuals’ fitness decreases with increasing challenge. As such, baseline glucocorticoid levels should be negatively related to fitness. In other words, the Cort-Fitness hypothesis suggests that high baseline glucocorticoids indicates that individuals are encountering challenging environmental conditions and have a low fitness compared to con-specifics in higher quality conditions. Environmental challenges can be predictable, semi-predictable, or unpredictable and the predictability and intensity of events can be independent. For instance, predation can be considered an unpredictable event, but seeing a predator and actually surviving a predation attack are very different in terms of the intensity of the predation event. On the other hand, breeding is a predictable event for birds that can be less intense only if the bird has some experience, even when assuming perfect resource conditions, which adds yet another layer of contributing factors, that is the life history of individuals. In breeding, the life history may go hand in hand with the age of individuals, but that is not the case for other events, as individuals experiencing similar challenging conditions show a variation in cort response levels based on their life history and what they experienced prior to the event (Satterthwaite et al., 2010; Zimmer et al., 2013; Bowers et al., 2016; Angelier et al., 2007).

## 2 Cort Fitness Mathematical Model

Mathematical modeling can be used to explore the effects of environmental challenges on biological systems (Dingemanse et al., 2004; Armbruster and Reed, 2005). For the Cort-Fitness hypothesis, environmental challenges can affect both the level of baseline cort and the individual’s fitness. Here, we investigate how birds’ cort level and fitness covary through a mathematical model that incorporates environmental challenges. We assume that all birds are living under the same environmental conditions and we formulate our mathematical equations based on birds’ characteristics. We make three structural assumptions as we formulate the model:

1. Three levels of predictability and intensity in terms of the challenges the animal faces: predictable, semi-predictable, unpredictable (Figure 3). We define predictable events as those that are repeatable (seasonal/periodic), can be long term such as breeding, reproduction, parental care, and migration and tend to be low intensity and chronic. Semi-predictable events occur stochastically, but follow a predictable temporal and spatial distribution, such as storms and temperature changes as well as as food availability and are often more intense. Unpredictable events are outliers from the baseline distributions of stochastic event and can be short term such as predation, or even the presence of predators and are of high intensity and acuteness.
2. Three groups of individuals based on their life history, specifically reproductive output: low, medium, and high fecundity.
3. Intensity of environmental challenges, cort level, and fitness vary along on a scale representing a minimum threshold to maximum level.

We follow a similar approach to (Johnson and Mangel, 2006; Mylius and Diekmann, 1995; Jones, 2011) which use the Euler-Lotka (E-L) model to derive the extrinsic rate of natural increase, *r*, of a population (our chosen fitness metric) and explore how life history choices affect fitness in simple organisms. The E-L model requires system specific mortality and birth functions. We introduce the chosen functions in the next few sections before combining them with the E-L equation and deriving *r*.

### 2.1 Cort Response and Reproductive Success

Brood value is a consequence of a strategic investment in reproduction. Brood value increases the population fitness *r* through maximizing the reproductive success of individuals *b_x_*. We assume that the birth rate function *b_x_* depends on cort level *ρ* (when the cort level is higher than a threshold) as well as on the minimum breeding age *T*, below which birds do not breed, but above which, in the simplest formulation of this model, the fecundity is constant. We will later consider relaxing this assumption since younger birds (close to *T*) will distribute their reproduction efforts among many attempts (i.e their brood value is low) while older birds will focus on fewer attempts to maximize their fitness (i.e. their brood value is high) (Satterthwaite et al., 2010; Bokony et al., 2009; Bouwhuis et al., 2009).

To explore the impact of differential fecundity and response to stress can impact fitness, we begin by defining three groups, representing 3 subpopulations. The groups are defined as follows:

- Group 1: low fecundity, high stress response.
- Group 2: high fecundity, low stress response.
- Group 3: medium fecundity, medium stress response.

These subpopulations may represent groups of individuals with different genotypes or differences in habitat quality. We assume that the age-specific fecundity of each group, *b_x,i_*(*ρ*), is described by two sets of equations. First, birds reproduce with constant fecundity, *f_i_*(*ρ*) that depends on cort levels, *ρ*, from when they become sexually mature at time *T*:

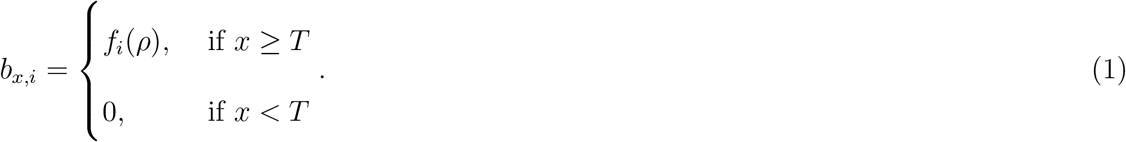

Here the reproductive success of the *i*^th^ group is denoted by *f_i_*(*ρ*). It represents birds’ reproductive success as their cort level changes and is defined as

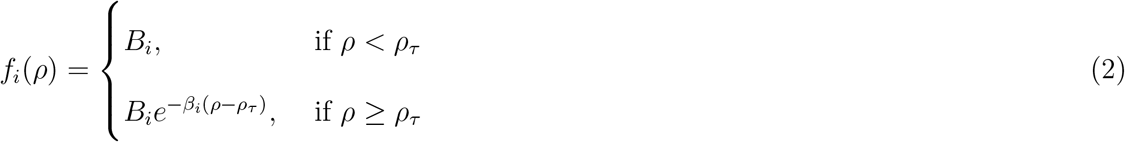

where *ρ_τ_* is a threshold cort level, *B_i_* is the birth rate of birds in group *i*, and *e*^−*β_i_*(*ρ*−*ρτ*)^ is the group-specific reduction in fecundity due to high cort levels (*ρ* ≥ *ρ_τ_*) for birds in group *i*. In figure 2 we show the reproductive performance across cort levels implied by these equations. The curves show that the reproduction rate is constant for resting state cort level (i.e. *ρ* ≤ *ρ_τ_*), but as birds become stressed, cort levels increase and the reproduction rate decreases. Other possible situations exist. For instance, in some situations exhibiting a hormonal stress response is favorable to individuals. They manage to adapt and perform better under stressful situations as described by the Cort-Adaptation hypothesis (Bonier et al., 2009a; Escribano-Avila et al., 2013). Our model does not address this case.

**Figure 2:**
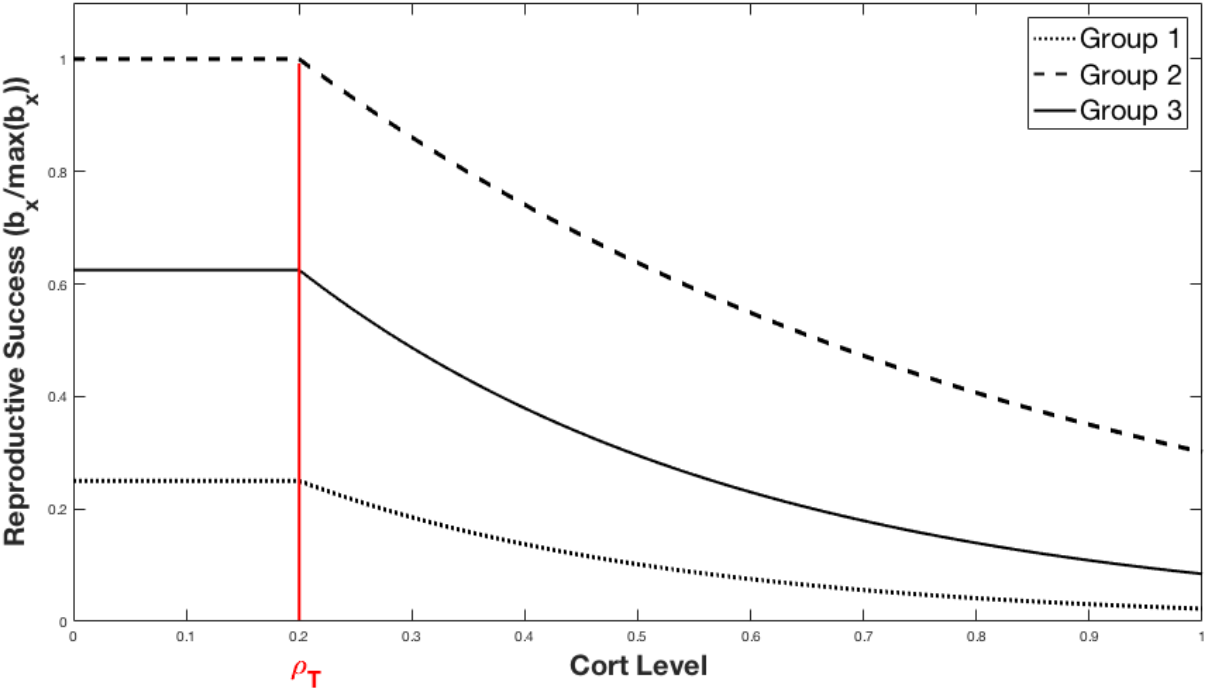
The reproductive rate function *b_x_* varies with cort level. The function starts as a constant and drops as cort level surpasses a threshold *ρ*_τ_. Birds’ reproductive success for each group category is negatively correlated with the stress level in the case of an unpredicted event. Parameter values for each group are presented in the appendix.

### 2.2 Environmental Challenges and Cort

Now that we have defined the proposed relationship between reproductive rate and cort levels, we link the cort levels back to the kind of environmental challenges (EC) experienced by birds. To model the non-linear relationship between cort levels, *ρ*, and environmental challenges (EC), we use a Hill function which was first introduced to describe the binding of macro-molecules (Hill, 1910). Here, we use it to describe the possible patterns of cort levels in response to a challenging environment. Based on previous studies (Lanctot et al., 2003; Romero and Wikelski, 2001; Jaatinen et al., 2013; Bonier et al., 2009b; Romero et al., 2000a) we assume a positive dependence, i.e. that cort levels increase with environmental challenges, but saturate. The dependence of cort on the EC is given by

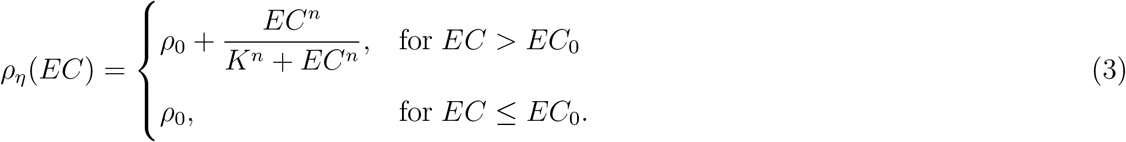

where *K* corresponds to the half-saturation constant, *n* is the Hill coefficient (*n* ≥ 1 for positive dependence), and *ρ*_0_ is a threshold cort level. This formulation explicitly assumes that challenges of low intensity are tolerable and so do not raise cort levels above the minimum, *ρ*_0_. Note that the thresholds *ρ*_0_ and *EC*_0_ are system specific.

We further assume that environmental challenges vary in terms of predictability as well as intensity. We consider three predictability levels (Figure 3 a), denoted by *η*, corresponding to different values of the parameters in equation (2.2): predictable (*η* = 1); semi-predictable (*η* = 2); and unpredictable (*η* = 3). These environmental challenges are further scaled for intensity from low (*EC* = 0) to high (*EC* = 1).

**Figure 3:**
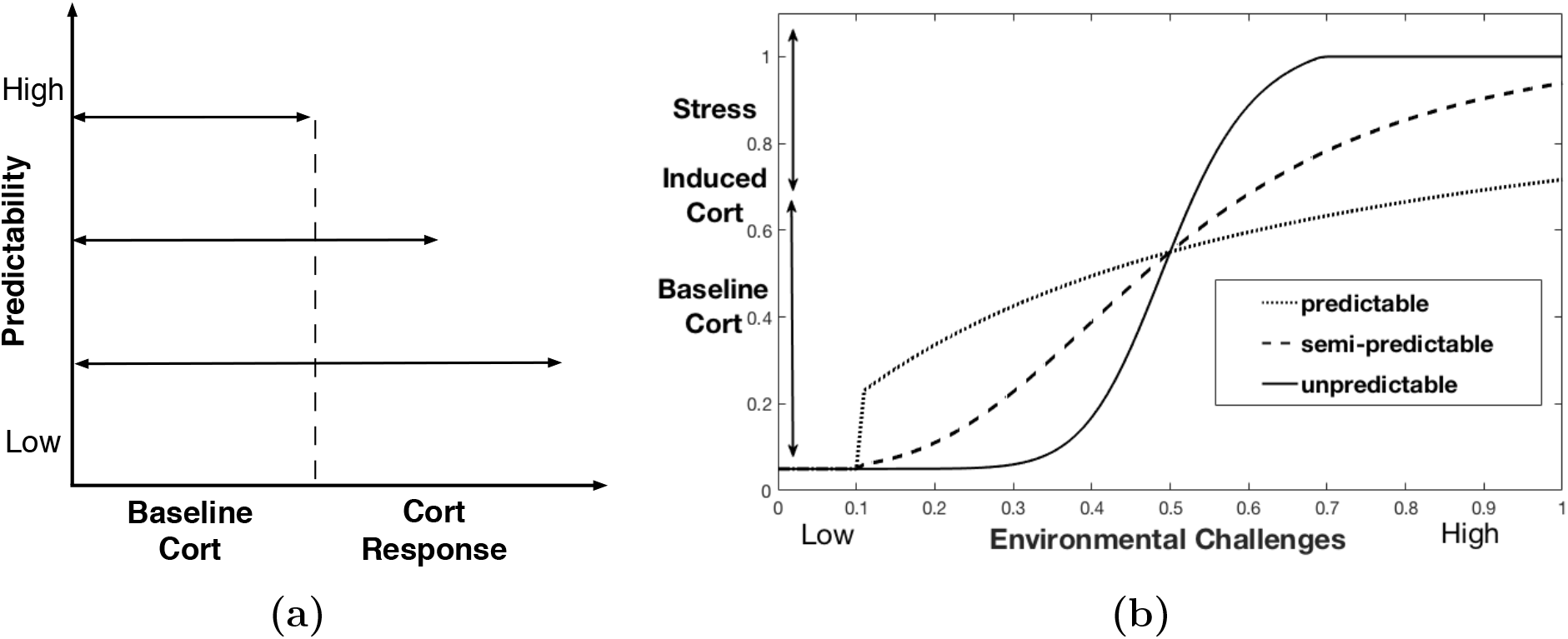
(a) An illustration of cort level variation within the baseline range and as a stress response based on the predictability of environmental challenges. (b) The function *ρ_η_* that describes the cort response to environmental challenges (EC) as they vary in predictability and intensity.

In Figure 3 we show the resulting cort response curves corresponding to the three environmental challenge scenarios as described by the respective Hill function for each predictability level. If birds experience predictable challenges, such as breeding, birds will anticipate the event and their cort level can build up while remaining within the baseline range. For semi and unpredictable scenarios, there is no prior anticipation of the challenging events and they are of higher intensity, so the level of cort increases dramatically as the intensity of the challenge increases. Galhardo et al. 2011 showed in an experiment a decreased stress response (lower/slower rise in cort) from male *cichlid* fish when the experienced stress event is predictable (Galhardo et al., 2011). Moreover, Romero et al. 2000 showed that within the breeding season, otherwise stressful weather events had less effect on the cort stress response in free-living birds. This suggests that birds prepare for the energetic demands of the breeding season, and events that would otherwise elicit dramatic changes in cort levels can be weathered. Predictable stressors that can be prepared for are no longer stressful (Romero et al., 2000b). Animals in general specifically birds, can predict storms and it alters their stress response in the same way. Storms are a common occurrence during the breeding season, and birds have developed ways of predicting when storms will occur and are able to prepare. This allows them to physiologically mediate stressful events to decrease the negative effects on reproduction. It provides support for the idea that a predictable event becomes less stressful and birds can increase their reproductive success by predicting these events (Breuner et al., 2013).

### 2.3 Fitness and Environmental Challenges

With the results in the previous two sections it is possible to examine how different assumptions in environmental challenges impact cort levels and then impact the fecundity of the birds. However, we seek to examine a more holistic measure of fitness, the intrinsic rate of natural increase of a population, *r*, which we can derive based on the Euler-Lotka (E-L) equation:

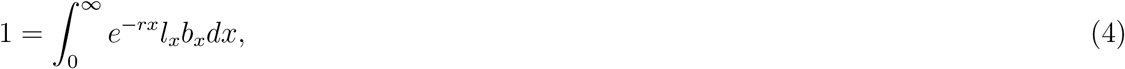

where *l_x_* is the probability that a female bird survives to age *x, b_x_* is the rate of reproduction of a female of age *x*, and *r* is the the intrinsic rate of natural increase, our chosen fitness measure. We assume that the survival probability for birds follows a type II function, that is an exponential decay curve (Demetrius, 1978), so that *l_x_* = *e^−mx^*, with m being the extrinsic mortality rate. With this assumption, the E-L equation for birds that reproduce at ages greater or equal to the minimum breeding age *T* is given by

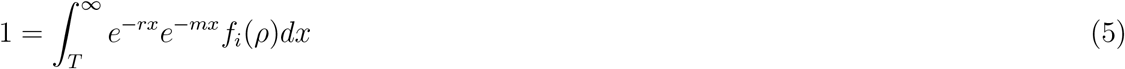

Integrating this equation we obtain the following transcendental equation for *r*:

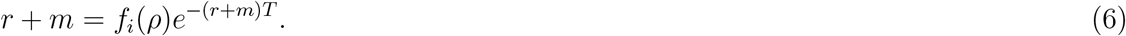

We then substitute the cort function *f_i_*(*ρ*) and obtain two cases:

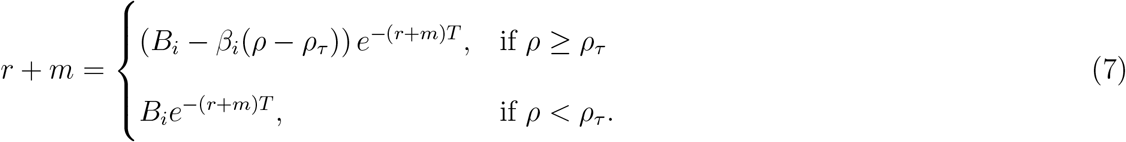

These equations may be solved numerically to obtain *r*. Further details on the derivation of *r* and numerical solutions can be found in the appendix.

We can now use two fitness metrics, *b_x_* and *r*, to investigate the effect of intensity levels for an unpredictable event on birds. In Figure 4 we show that the reproductive rate *b_x_* has a similar pattern across all bird groups that experience unpredictable events. All groups start with a constant reproductive rate assigned to their group, then, as the intensity of environmental challenges increases the reproductive success decays at different rates for each group, corresponding to the groups’ tolerance. We notice that for groups 1 and 3 (the low reproduction/high stress response and medium reproduction/medium stress response, respectively), reproductive success approaches zero. As for group 2 (high reproduction and low stress response), we see that the reproductive rate decreases but does not approach zero.

**Figure 4:**
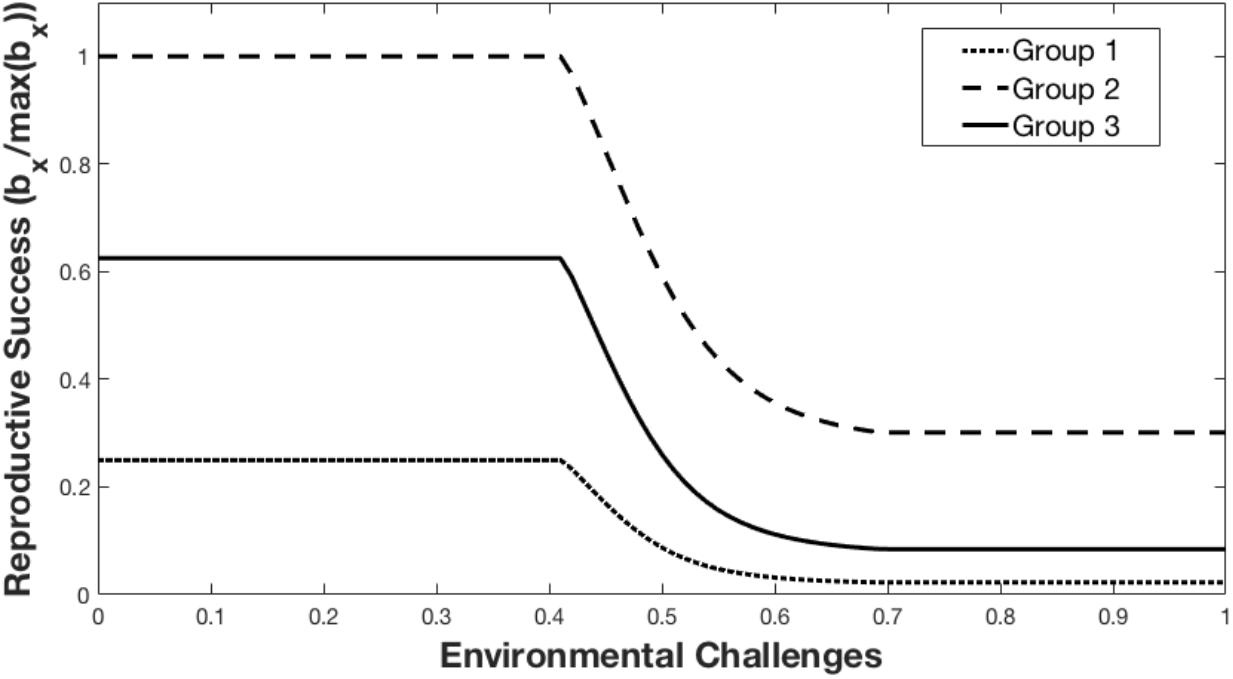
Reproductive success *b_x_* for birds’ three groups varying as we increase intensity for an unpredictable event.

In Figure 5 we show the response of our second fitness metric, the intrinsic rate of natural increase *r*, as a function of environmental challenge for the three groups. Here a positive value of *r* indicates that the population is increasing, while a negative value is an indication that the population is declining. Thus, *r* for each group category gives an estimate of the relative performance of bird populations composed of members that share the same characteristics. Given the same assumptions as in Figure 4, i.e an unpredictable event, in Figure 5 we show that *r* decreases as the intensity of challenging events increases, due to the incorporation of the EC dependent birth rate for each group. Moreover, we see that for a population of birds with group 1 characteristics, *r* drops below zero as EC intensity to about 50% of the maximal intensity (lowest curve) – that is this population is expected to decrease if the intensity of environmental challenges is high. Group 3 (our medium characteristic group) shows a similar pattern, although this group can handle higher intensity EC. In contrast, the population with group 2 characteristics is able to maintain positive population growth (*r* > 0) even at the highest intensities.

**Figure 5:**
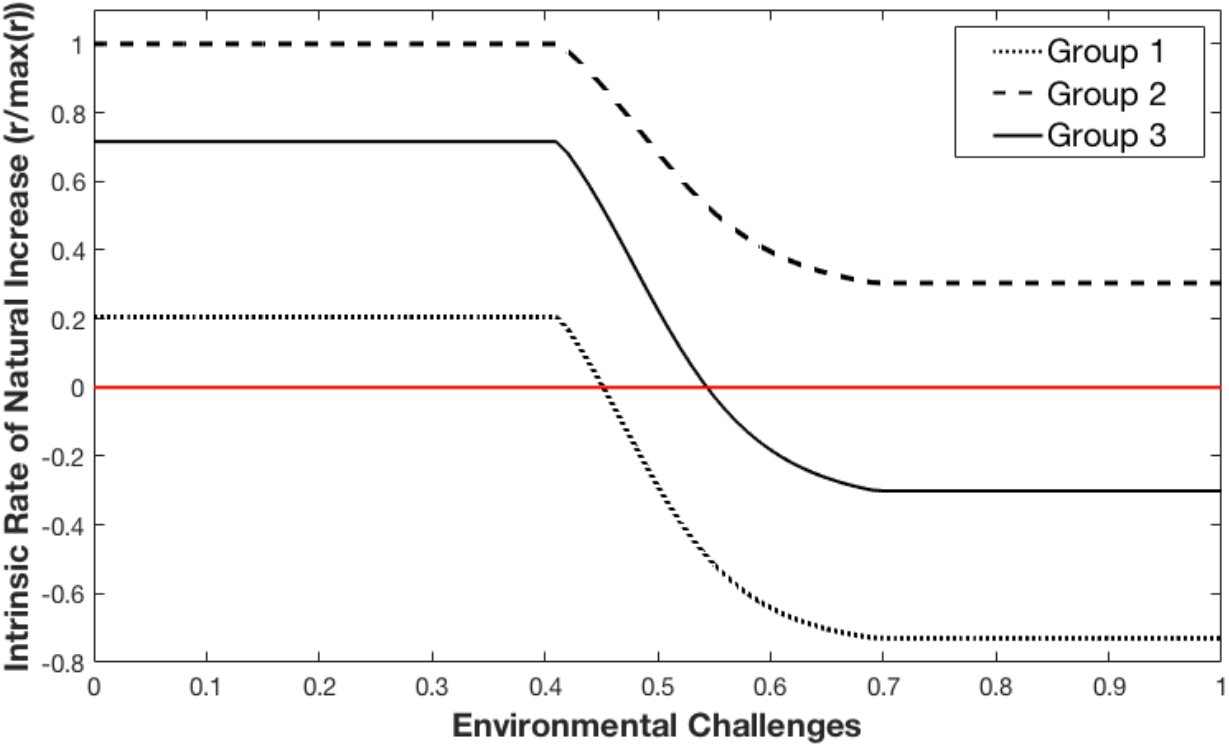
The intrinsic rate of natural increase r for birds’ three groups varying as we increase intensity for an unpredictable event.

Similarly, Figure 6 shows the intrinsic rate of natural increase as a function of cort level. Each group maintains a constant intrinsic rate when the cort level is under a threshold *c_T_*. Beyond the cort threshold, the intrinsic rate of natural increase shows a negative correlation with high cort levels. Overall, birds with group 2 characteristics perform better than birds in group 1 and 3. Patterns are qualitatively similar for the predictable and semi-predictable scenarios (see appendix).

**Figure 6:**
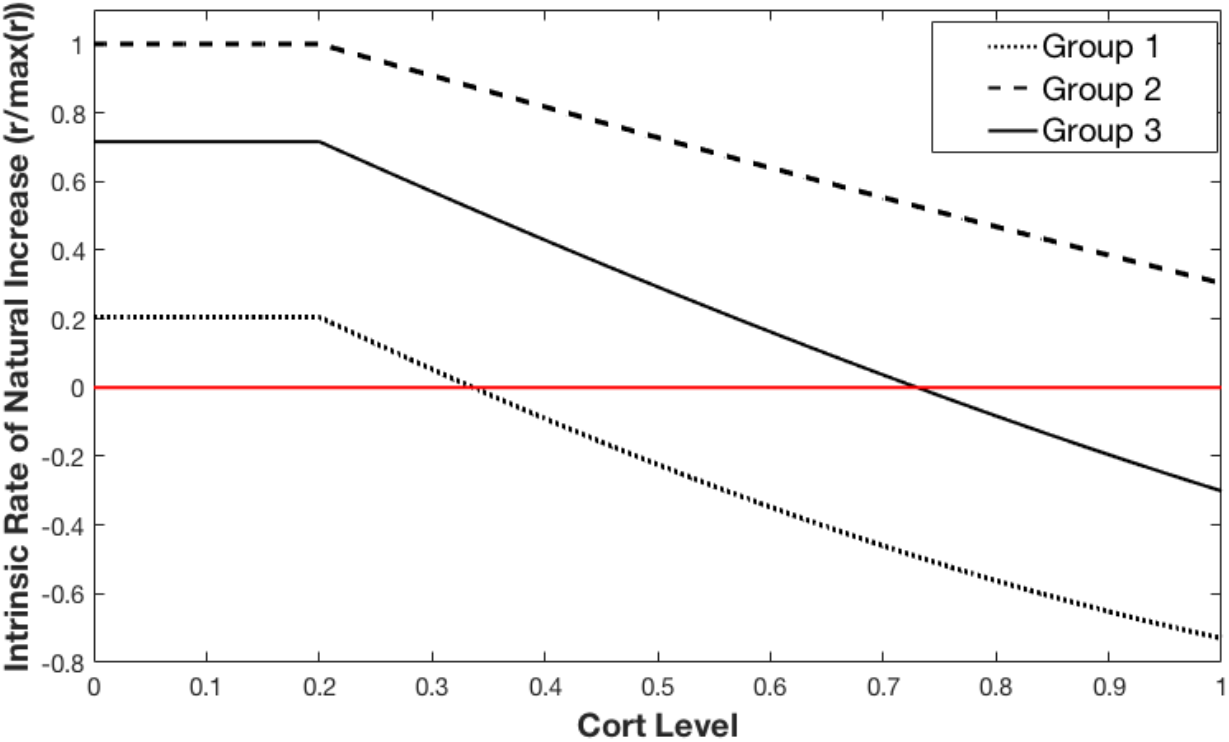
The intrinsic rate of natural increase value versus cort level.

## 3 Discussion

The relationship between environmental challenges, glucocorticoid hormones, and fitness is very complicated. So far, it has been primarily investigated through observational studies, but there are many limitations that come with that such as the non-repeatability of these studies, the non-linear dependence between the three components, the choice of fitness metric, as well as individuals’ own life history that shapes how they cope with stress. Here, we propose a mathematical model that explores different environmental challenging scenarios and how they affect cort and fitness. The model assumptions include: (1) non-linear dependence between environmental challenges, cort, and fitness; (2) two fitness metrics, namely, reproductive success and the intrinsic rate of natural increase; (3) environmental challenges varying in predictability and intensity. The model is build based on the Cort-Fitness hypothesis assumptions but can be generalized to also address cort as being a positive driver for short lived species to increase their fitness (e.g., Cort-Adaptation hypothesis). We first divided the population into three groups that differ in baseline reproductive output and stress response. Then we looked at how individuals from each group respond to stress caused by an unpredictable challenging event (see Figure 2). Next, we explored the effect of environmental challenges on cort by allowing both intensity and predictability to vary. Figure 3 provides the three possible scenarios of cort response to a predictable, semi-predictable, and unpredictable environmental challenges that go from low intensity to high. These curves can be used differently depending on the event we are interested in exploring. For instance, the curve for the unpredictable event was used to obtain results in Figure 4 and Figure 5. The choice of two fitness metrics allowed the model to extend the information from the individual to the population level which can be useful in conservation endocrinology where cort is used as a proxy to discuss individuals trade off strategy under challenging conditions (Satterthwaite et al., 2010; Jaatinen et al., 2013; Satterthwaite et al., 2012). Moreover, we can use the model to track individuals’ progress as they acquire more experience and their brood value increases. Figure 7 shows the fitness measure across the 3 groups. We could imagine that, instead of the separate groups as we showed in our analysis here, that groups could represent different subpopulations utilizing different environments or with different phenotypes. The model could even potentially be extended to allow the groups to represent differences of a population across life stages. For example, young individuals could start in group 1 with limited experience and low breeding success, therefore a low fitness for the (sub)population. Then as they gain experience, improve their coping mechanism, they increase their breeding success, and the fitness of the subpopulation gets higher until it drops as they get older and senesce. The model in its current form does not give the explicit formulation for this age dependent case (however, see the appendix for this age dependent case) our simple model can give an example of how observed patterns of fitness as a function of cort and EC can depend on the make-up of the specific population being observed. Because of this sort of variation between individuals and populations, exploring the Cort-Fitness hypothesis can be very challenging. While previous investigators have developed models to help understand the mechanisms, effects, and variation in hormonal stress responses (e.g., the allostasis model ((McEwen and Wingfield, 2003)) and reactive scope model ((Romero et al., 2009))), they have not been mathematically based.we envision that our mathematical model can be used as a first step to explore specific hypotheses about the effects of timing and intensity of challenges on cort levels and fitness and ultimately develop a better understanding the complicated patterns that have been observed currently.

**Figure 7:**
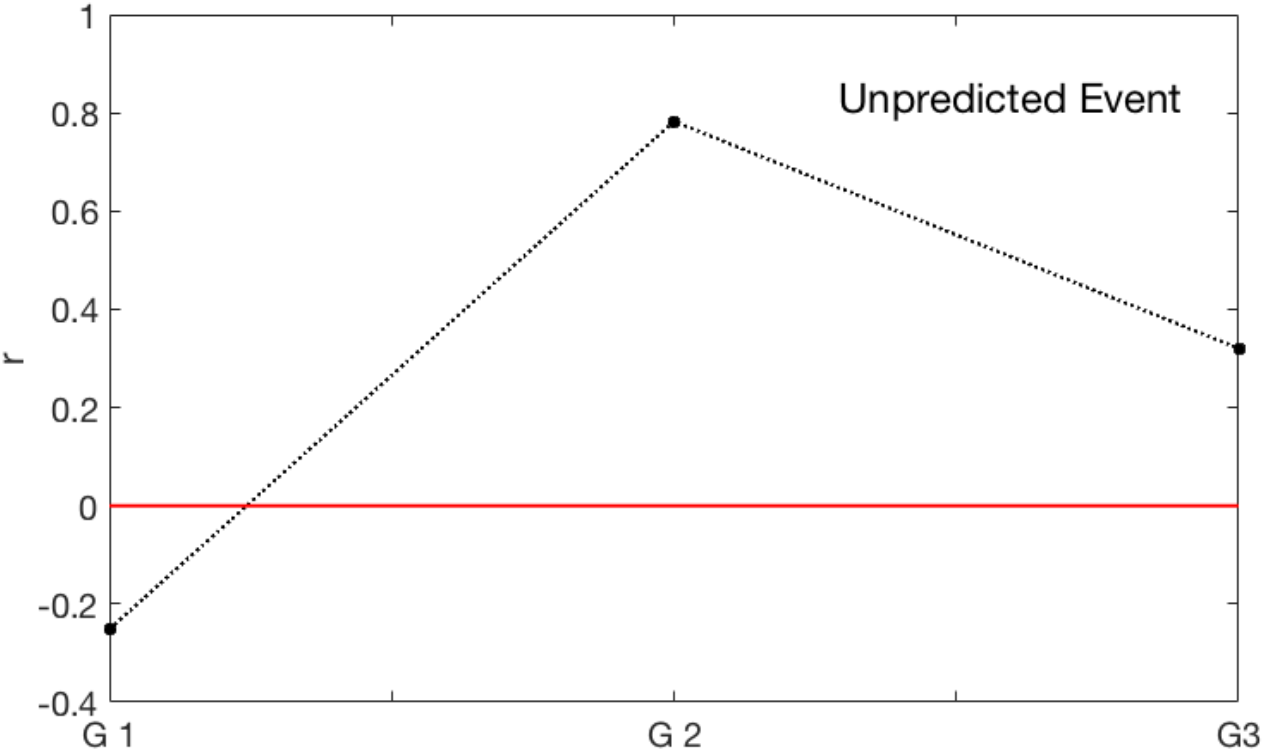
The intrinsic rate of natural increase value for the three groups at a fixed level of EC for an unpredictable event.

## Acknowledgment

We thank Frances Bonier for invaluable comments on early drafts.

## Authors’ Contributions

F.E.M., I.T.M., and L.R.J. designed the study. F.E.M. and L.R.J. performed the analyses. F.E.M, S.J.L., and I.T.M. parameterized the model. F.E.M. wrote the paper. All authors contributed to revising and editing the paper.

## A Appendix

### A.1 More Results

Here we have results for all scenarios proposed by our model. Figure 6(a) shows both fitness metrics, the reproductive success *f* and the intrinsic rate of natural increase *r* varying with cort level ranging from baseline cort to a stress response. For each of the three population groups (group 1, 2, and 3) we investigate three different challenging event, predictable, semi-predictable, and unpredictable. While Figure 8(b) shows the fitness metrics varying with environmental challenges intensity rather than cort level. However, since cort and challenging events are positively correlated, we observe a similar trend. Overall, the results show that individuals in group 2 have a better performance in all challenging scenarios when compared to groups 3 and 1. Which is what we would expect given these individuals experience and brood value. We notice that in extreme challenging scenarios associated with very high cort level, individuals’ reproductive success becomes very low. This leads to low values of the intrinsic rate of natural increase that is associated with a decay in the overall population.

**Figure 8:**
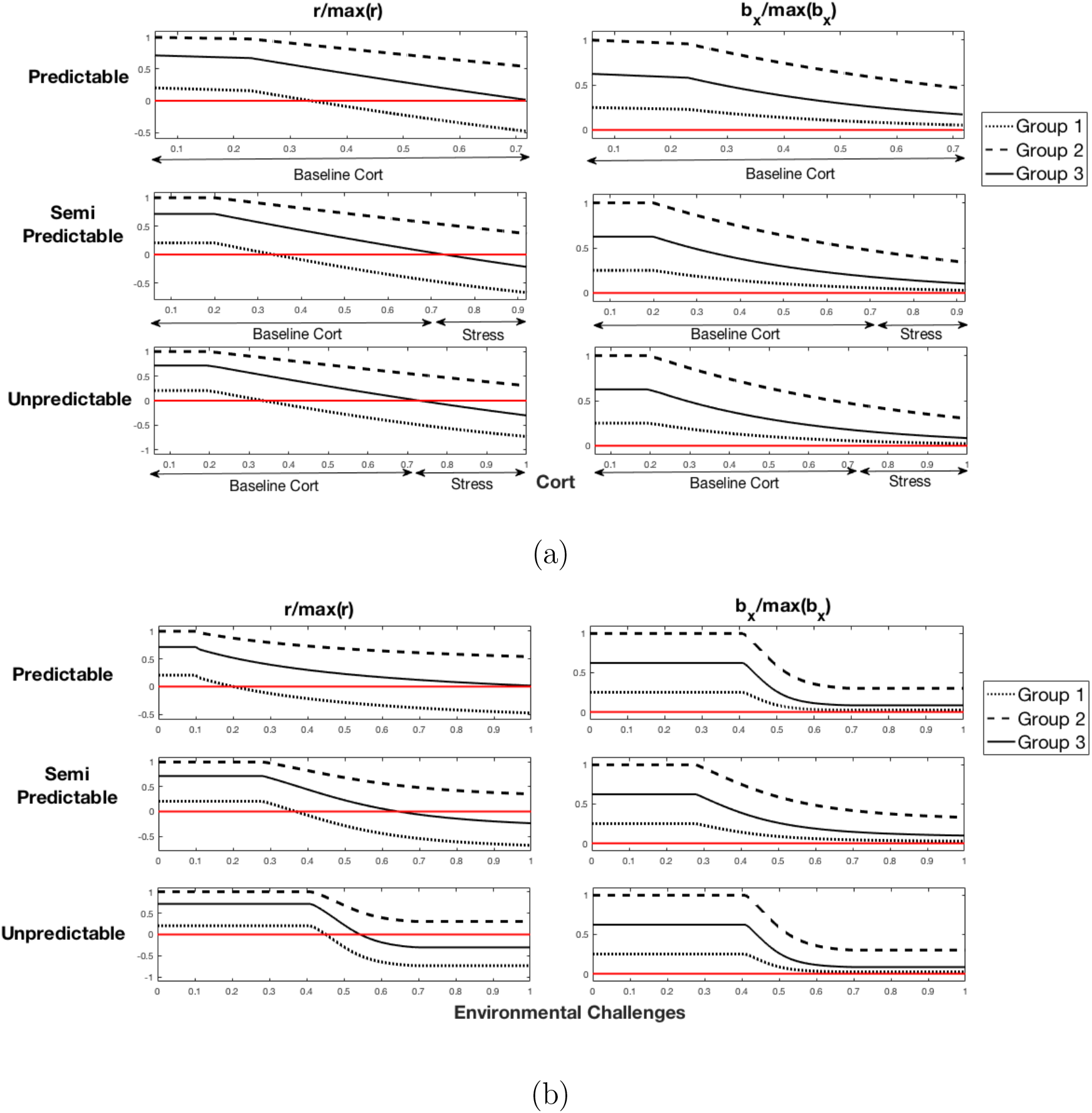
Fitness of three bird groups varying with cort level and challenges intensity under three different challenging events scenarios.

### A.2 Deriving the intrinsic rate of natural increase

Here we only consider birds capable of breeding (i.e. with age ≥ T). To derive the intrinsic rate of natural increase from the Euler-Lotka equation we proceed as follows:

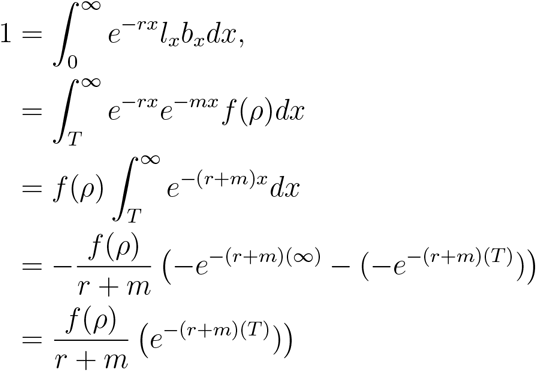

which yields

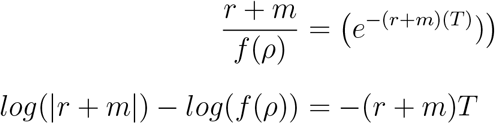

Hence,

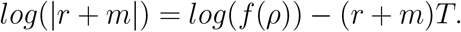

### A.3 Numerical Solution and Parameter values

To solve for r we use Newton-Raphson numerical method ((Walston, 1968; Ypma, 1995)). All equations are solved using Matlab and the code can be found in our git repository https://github.com/fadzEM/Cort_Fitness_Model

The parameters used in the model and their description are given in Table 1.

**Table 1:** Parameters used in the model.

| Parameter | Value | Description | Unit |
| --- | --- | --- | --- |
| n | 1 | Hill coefficient | no unit |
| K | 5 | Hill half saturation constant | EC |
| T | 2 | age at which birds start breeding | time |
| $\beta_1$ | 3 | decay rate for group 1 | 1/time |
| $\beta_2$ | 1.5 | decay rate for group 2 | 1/time |
| $\beta_3$ | 2.5 | decay rate for group 3 | 1/time |
| B | 15 | constant growth rate for bird population is $\alpha \times B$ | number of fledglings |
| $\alpha_1$ | .2 | $\alpha$ for group 1 | 1/time |
| $\alpha_2$ | .8 | $\alpha$ for group 2 | 1/time |
| $\alpha_3$ | .5 | $\alpha$ for group 3 | 1/time |
| $c_T$ | .2 | cort threshold | cort |
| $c_0$ | .05 | cort minimal value | cort |
| m | .6 | birds natural mortality rate | 1/time |

### A.4 Age Dependent variation in fecundity and stress responses

When birds complete their first molt and reach age *T*, they begin mating and participate in breeding seasons. Figure 9 shows an illustration of birds’ success of reproduction as they age, defined by the number of fledglings they produce. Birds in group 1 start their first breeding attempts at age *T* and will have a few failed breeding attempts before reaching a mature age which allows successful breeding. Birds in group 2, i.e. with age a bit greater than *T*, have enough experience to perform multiple successful breeding seasons and tolerate higher cort levels better than they did when in group 1. Eventually they get older and move to group 3 and have reduced reproduction and are less tolerant to high cort levels.

**Figure 9:**
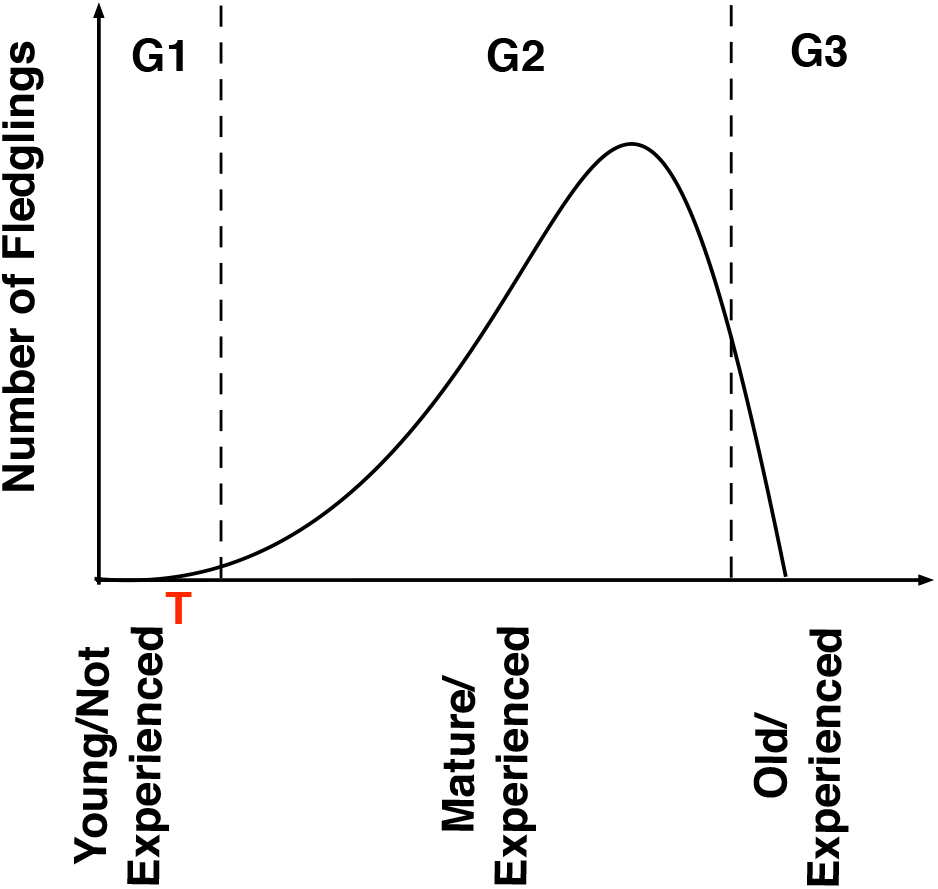
We assume that the birds’ population is composed of three groups based on individuals breeding experience. As illustrated in the graph, birds under age T do not succeed in breeding (*G*_1_). As they age, they develop more experience (*G*_2_), and become more successful in breeding. When they get older they gradually become less successful (*G*_3_).

We seek to capture the age dependent variation by extending our model. First we define the times that mark the transitions between groups

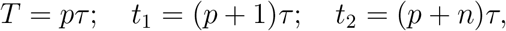

so that *τ* is the time birds spend in group 1, (*n* − 1)*τ* is the time spent in group 2, and the remaining time spent in group 3 is what is left until the birds die. In this case, the E-L equation for the overall population is given by

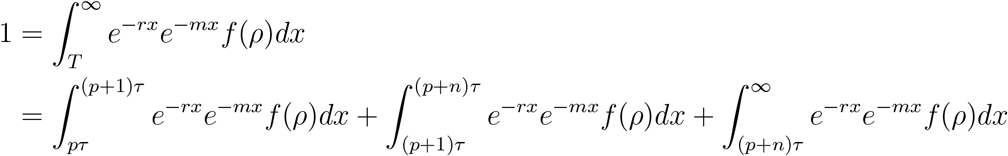

integrating this and simplifying we obtain:

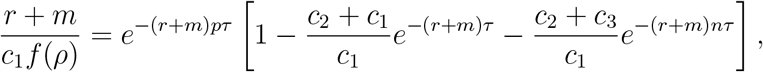

where *c*_1_, *c*_2_, *c*_3_ are constants of integration. Similarly to the simpler version of the model, we can use Newton-Raphson method to approximate the intrinsic rate of natural increase numerically. This method is species specific as it requires the knowledge of the amount of time individuals spend in each group.

